# Stereo-anomaly is found more frequently in tasks that require discrimination between depths

**DOI:** 10.1101/2023.11.02.564189

**Authors:** Alex S Baldwin, Seung Hyun Min, Sara Alarcon Carrillo, Zili Wang, Ziyun Cheng, Jiawei Zhou, Robert F Hess

**Affiliations:** McGill Vision Research, Department of Ophthalmology & Visual Sciences, McGill University, Montreal, Quebec, Canada; The Eye Hospital, School of Ophthalmology and Optometry, Wenzhou Medical University, Wenzhou, Zhejiang, China

**Keywords:** binocular vision, depth, disparity, individual differences, stereo vision, stereoacuity, stereopsis

## Abstract

Within the population of humans with otherwise normal vision, there exists some proportion whose ability to perceive depth from binocular disparity is poor or absent. The prevalence of this “stereoanomaly” has been investigated in previous studies, some finding the proportion to be as small as 2%, others finding it to be as great as 30%. In this study, we set out to investigate the possible reason for the wide range of results found in these studies. We used a digital stereoacuity measurement tool that could measure performance in tasks requiring either the detection of disparity or the discrimination of the sign of disparity. The stimulus design was otherwise similar between the two tasks. In a cohort of 228 participants, we found that 98% were able to consistently perform the detection task. In contrast, only 69% consistently performed the discrimination task. The 31% of participants who had difficulty with the discrimination task could further be divided into 17% who were consistently unable to perform the task (seeming to behave at chance), and 14% who showed some ability to perform the task. We propose that the greater prevalence of stereo-anomaly is revealed when tasks require the judgement of the direction of disparity.

## 1 Introduction

In human vision, the horizontal separation of the two eyes can be exploited to determine the relative distances to objects in the outside world. Fixation at a point in 3D space converges the eyes and establishes a curved surface of zero retinal disparity (the horopter). Objects closer or further away from the horopter will have retinal images projected at horizontally shifted locations in the two eyes (disparity). When the objects are closer the disparity is said to be crossed. Objects further than the horopter are said to be in uncrossed disparity. Humans can show exquisite sensitivity to binocular disparity when making judgements of the relative depth of objects.

Within the human population, some individuals lack the ability to determine depth from disparity (stereo-blindness) or else show an impairment in that ability (stereo-anomaly). In some cases, this is due to an identified disease (such as amblyopia) but in other cases the ability can be limited with no apparent cause. There is controversy over how common this latter condition is. Some studies have found stereo-anomaly in up to 30% of participants (Richards, 1970; Hess et al., 2015). Other studies have found these cases to be much rarer, affecting as few as 1-2% of individuals (Newhouse and Uttal, 1982; Coutant and Westheimer, 1993; Birch et al., 2008; Bohr and Read, 2013).

More recent studies have implemented stereo testing on handheld digital tablet devices; this approach has been proposed as a convenient clinical tool. One study that used a random-dot stimulus displayed on a tablet found that about 30% of the population with otherwise normal vision were up to ten times worse in their stereo sensitivity (Hess et al., 2016). This was surprising, as the widely-used Randot clinical stereo test exhibits a very narrow distribution of stereo sensitivity in that population (Birch et al., 2008). Although these two approaches to clinical stereo measurement are different in several ways, one possible explanation of this difference is particularly interesting: stereo deficits may be specific to tasks where the polarity of the depth must be discriminated (Landers and Cormack, 1997; Hess et al., 2019). These stereo-anomalies would not be found with tests such as the Randot, which rely simply on detecting shapes defined by disparity. In the present study, we compare results from disparity detection and disparity discrimination tasks using otherwise equivalent stimuli in the same individuals. Our goal is to determine the basis of the previously reported stereo deficit in individuals with otherwise normal vision.

## 2 Methods

### 2.1 Participants

We tested 238 participants (aged 17-36, 150 female). All subjects had normal vision or were corrected-to-normal with prescribed optical correction (above 20/20). Subjects reporting any visual disorders (cataracts, glaucoma, or amblyopia) were excluded from the study. Written informed consent was given by all subjects. All testing was performed in accordance with the Declaration of Helsinki and approved by the Research Ethics Board of the McGill University Health Centre and Wenzhou Medical University.

### 2.2 Apparatus

Testing was carried out using two stereoacuity apps developed at McGill Vision Research (Hess and Baldwin, 2020). These are variations on the Baldwin-Hess stereo test design that has been used in a number of previous studies (Tittes et al., 2019; Webber et al., 2019; Alarcon Carrillo et al., 2020; Atchison et al., 2020; Webber et al., 2020; Alarcon Carrillo et al., 2023). These were installed on a 5th generation Apple iPad mini (model A2133; Apple Inc, Cupertino, CA). Testing was performed at a viewing distance of 32 cm (maintained with the aid of a length of string attached to the tablet). At this distance, the display had 72 pixels per degree of visual angle. The screen brightness was set to about half the maximum level, approximately 150 cd/m^2^. The iPad’s automatic brightness control feature was turned off.

### 2.3 Stimuli

The stimuli in the detection and discrimination tasks were generated using methods similar to those reported in our previous studies (Tittes et al., 2019; Alarcon Carrillo et al., 2020, 2023). These studies share the use of stimuli generated from spatially-bandpass (peak spatial frequency 0.4 c/deg, spatial frequency bandwidth *±*2.2 octaves) isotropic log-Gabors which are composed into a field of “fuzzy” dots on a grey background. The current study uses a mixture of dots with dark and light centres, after Alarcon Carrillo et al. (2020). They were placed with an average dot-to-dot distance of 32 arc min. Stimuli were presented at a Michelson contrast of 80%.

Disparity was introduced into the stimuli by shifting the horizontal position of the dots. For the disparity detection task (Figure 1A) the dots falling within a wedge-shaped target zone (6 deg radius) were given a crossed disparity equal to half of the total stimulus disparity. The remaining “background” dots were given an equal amount of “uncrossed” disparity. The total stimulus disparity therefore is that between the foreground (wedge) and background dots. This has the benefit of applying the same positional shift to every dot in the stimulus, safeguarding against both binocular and monocular cues that might allow a participant without stereo vision to perform this task (O’shea and Blake, 1987; Serrano-Pedraza et al., 2016; Chopin et al., 2019).

**Figure 1.**
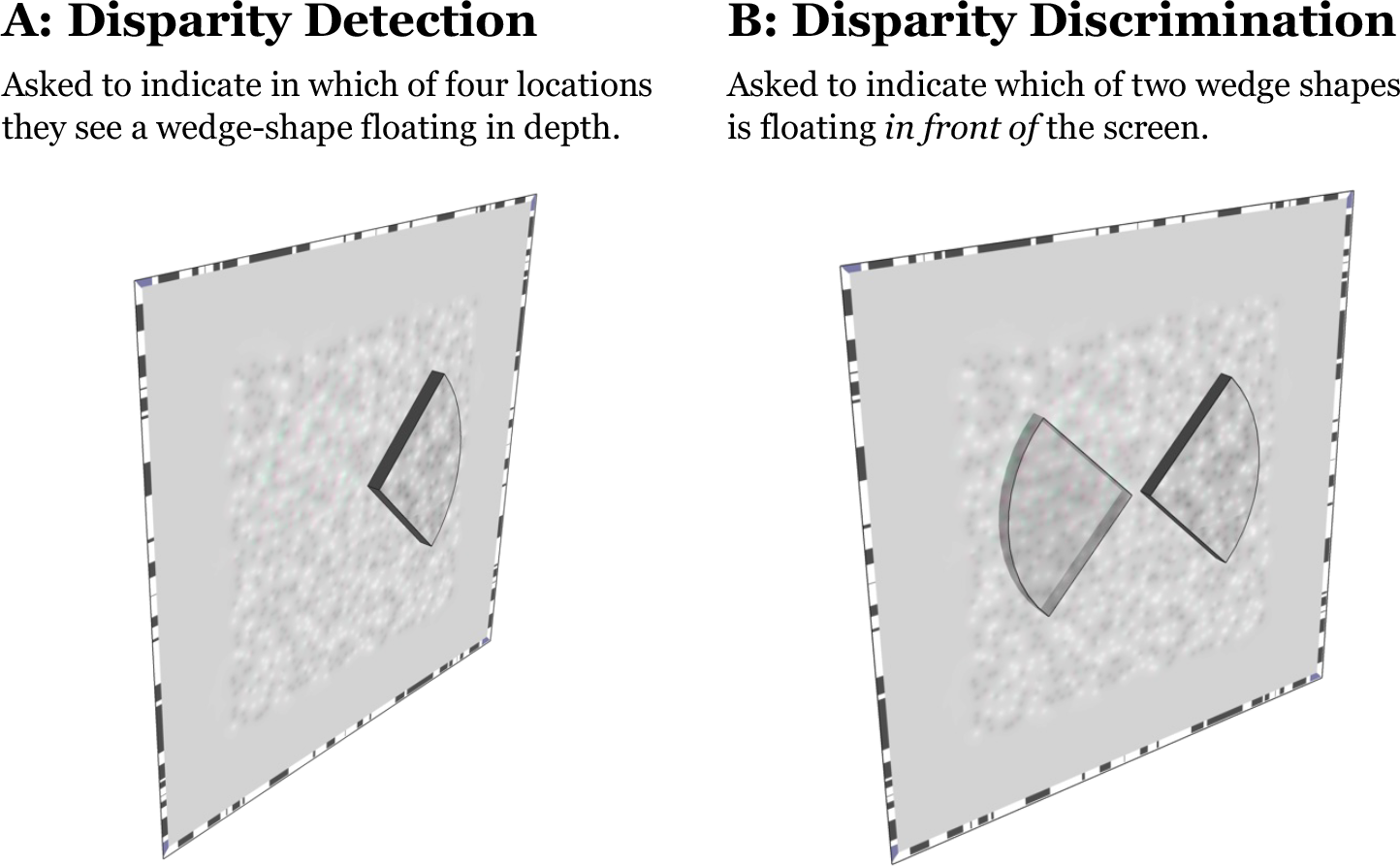
Diagrams showing stimulus design for the two measurements. Stimuli are rendered in a 3D projected view, with the wedges of disparity indicated. Panel **A** shows the design for the disparity detection task, in which there were four possible target locations. This example shows the target on the right. Panel **B** shows the design for the disparity discrimination task, in which there were two possible target locations. In this example, the wedge on the right has crossed disparity and so is the target.

The disparity discrimination task featured two wedge-shaped targets to which disparity was applied (Figure 1B), with the remaining dots being rendered at the plane of the display. The target wedge was rendered with crossed disparity, and the other wedge with uncrossed disparity.

### 2.4 Procedure

The disparity detection task was a four-alternative forced-choice task (target shown in the left, right, top, or bottom position), whereas the discrimination task was two-alternative forced-choice (target on the left or on the right). Disparity thresholds were obtained for both the detection and discrimination tasks. The disparities were selected on a trial-to-trial basis using a pair of interleaved two-down one-up staircase algorithms to sample the informative region of the psychometric function. The initial disparity was 1024 arc sec and the step size was a factor of two. Testing was performed indoors. Each participant performed the detection and discrimination tasks twice. The order of the four testing sessions was randomized to prevent bias.

### 2.5 Analysis

Psychometric functions were fitted using the Palamedes toolbox (Prins and Kingdom, 2018) in GNU Octave (Eaton et al., 2020). Further analysis and graph plotting was performed in Python (Python Software Foundation, Wilmington, DE) using the Matplotlib (Hunter, 2007) and NumPy (Harris et al., 2020) libraries. Intraclass Correlation Coefficients (ICC) were calculated using Pingouin (Vallat, 2018). Mann-Whitney U tests, Kruskal-Wallis H-tests, Wilcoxon signed-rank tests, and t-tests were performed using SciPy (Virtanen et al., 2020). Statsmodels (Seabold and Perktold, 2010) was used to calculate binomial confidence intervals.

In all analyses, we took the log_2_ of the disparity values and performed our fitting and analysis with this logarithmic scale. To fit the psychometric function, we used a Logistic function and obtained the threshold at 55.20% correct for the 4-alternative forced-choice detection task and 76.02% correct for the 2-alternative forced-choice discrimination task (corresponding to a d’ of 1 in both cases). The standard error associated with the threshold was obtained through non-parametric bootstrapping. We set criteria on the result of the psychometric function fitting to determine whether a valid result had been obtained. Invalid results were those with a large standard error (>2 log2 units) or those where the computed threshold was outside of the range measurable by the device (see Alarcon Carrillo et al., 2023). In this case that limit was 4,096 arc seconds (12 log_2_ units).

## 3 Results

The test-retest agreement for the two tasks is plotted in Figure 2. The data are divided according to the number of valid measurements obtained on each task. It is immediately apparent that there are many more invalid results with the discrimination task than with the detection task. This indicates a specific difficulty associated with the discrimination task. The test and retest distributions for each task are visualised as histograms in Appendix Figure A1. The median discrimination thresholds are 25-30% higher than the median detection thresholds for the test and retest measurements. The largest group obtained a valid measurement on both repetitions of both tasks (68%). The other two large groups were those who failed both repetitions (17%) or one repetition (14%) of the discrimination task. The proportion of invalid results found using the discrimination task decreases from 29% on the first (“test”) measurement to 21% on the second (“retest”) measurement.

**Figure 2.**
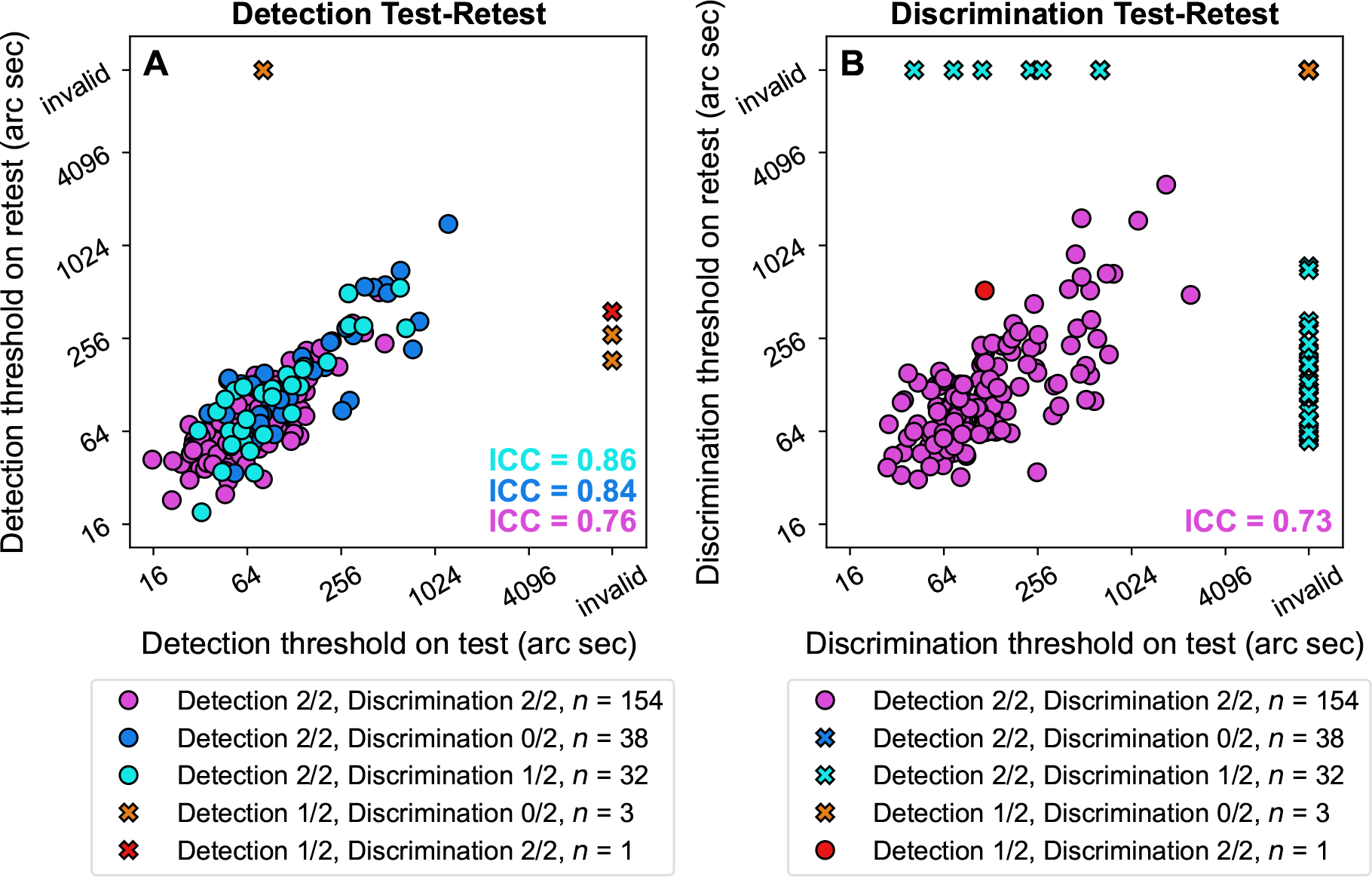
Test-retest scatter plots for the detection (**A**) and discrimination (**B**) tasks. Participants are divided according to whether two, one, or zero valid measurement results was obtained in each task. Intraclass Correlation Coefficient (ICC) scores were calculated using a two-way mixed-effects model for the groups with at least 30 participants. We use 2/2 to refer to participants where both test and retest measurements were valid, 1/2 to refer to cases where one of the two was valid, and 0/2 where neither were valid.

Test-retest agreement was assessed using two-way mixed-effects Intraclass Correlation Coefficient (ICC) scores. For the detection task all ICC scores fell in the “excellent” range. For discrimination, the ICC score was at the top of the “good” range (Cicchetti, 1994). These results show robust agreement between test and retest for each group of participants. A Bland-Altman analysis (Appendix Figure A2) revealed no significant bias; this means we do not find thresholds generally improve between the test and retest measurements.

We performed an additional analysis exploring the overall proportion-correct performance of the participants in the discrimination task, by collapsing over the different disparity levels (a method we previously used to look for residual stereopsis in amblyopic participants in Alarcon Carrillo et al., 2023). We calculated the 99% binomial confidence intervals of their overall proportion-correct. These were used to determine whether each participant appeared to be guessing (not significantly different from the 50% guess rate), or if there was evidence they performed better or worse than guessing (Appendix Figures A3-A4). We found that participants who failed both test and retest measurements were typically at chance performance. On the other hand, those who failed only one repetition tended to have performed above-chance in both repetitions. We can take this as evidence that the single valid result obtained from these individuals reflects an actual ability to perform the discrimination task, and is not due to chance guessing.

For clarity, the analysis from this point will not feature data from the small number of participants who failed any repetition of the detection task (four participants in total). We will return to these participants in our Discussion. The remaining participants are those from whom valid measurements were obtained in both repetitions of the detection task. We compared the performance of each participant on the two tasks. This is presented in Figure 3. The population was divided into three categories: those from whom we obtained a valid discrimination threshold on both the test and retest repetitions (pink, 154 participants), those who gave two invalid discrimination results (blue, 38 participants), and those from whom we obtained a single valid discrimination result (cyan, 32 participant). We used the mean of the test and retest measurements for participants where these were both valid. For participants where only one of the two was valid, we used the value from that valid measurement only.

**Figure 3.**
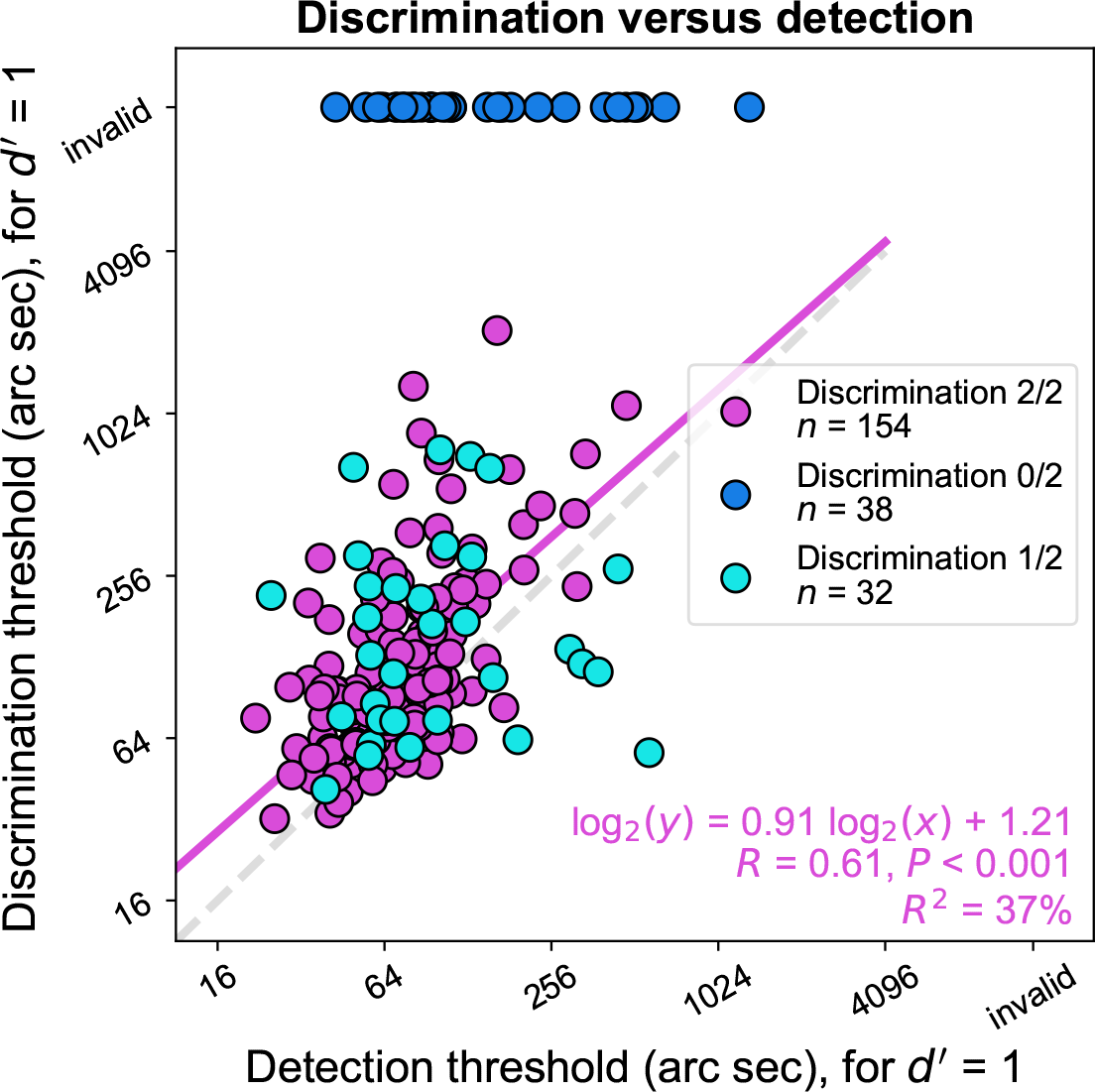
Scatter plot showing the relationship between disparity discrimination threshold and disparity detection threshold in the 224 participants from whom we obtained two valid detection thresholds. When thresholds were out of range the data points are placed at a disparity of 14,000 arc sec. A linear regression analysis was performed on the data from the individuals with normal stereo. This is shown by the pink line, and the statistics are reported in the bottom-right.

Among those who could perform both tasks (valid test and retest measurements for both detection and discrimination), we found a highly significant Pearson correlation between their detection and discrimination thresholds on double-log axes (*R* = 0.61, *P* < 0.001). The equation is shown in the bottom-right of Figure 3. The slope of the relationship was almost linear (0.91 on our double-log axes). We performed an additional analysis constraining the slope to be 1, in which case the offset value is 0.65. This means that the discrimination threshold for each participant was generally around 60% higher than their detection threshold.

The distributions of detection and discrimination thresholds from the three groups shown in Figure 3 are plotted as histograms in Figure 4. For the detection thresholds (left column), the median stereoacuity was best (69 arc sec) for those who obtained two valid results in the discrimination task. The highest median threshold (105 arc sec) was found in those who were unable to obtain a valid result in either repetition of the discrimination task. In the group who had a valid result in one of the two repetitions of the discrimination task the median threshold was intermediate (83 arc sec). This suggests that the variation in ability to perform the discrimination task is predictive of performance in the detection task.

**Figure 4.**
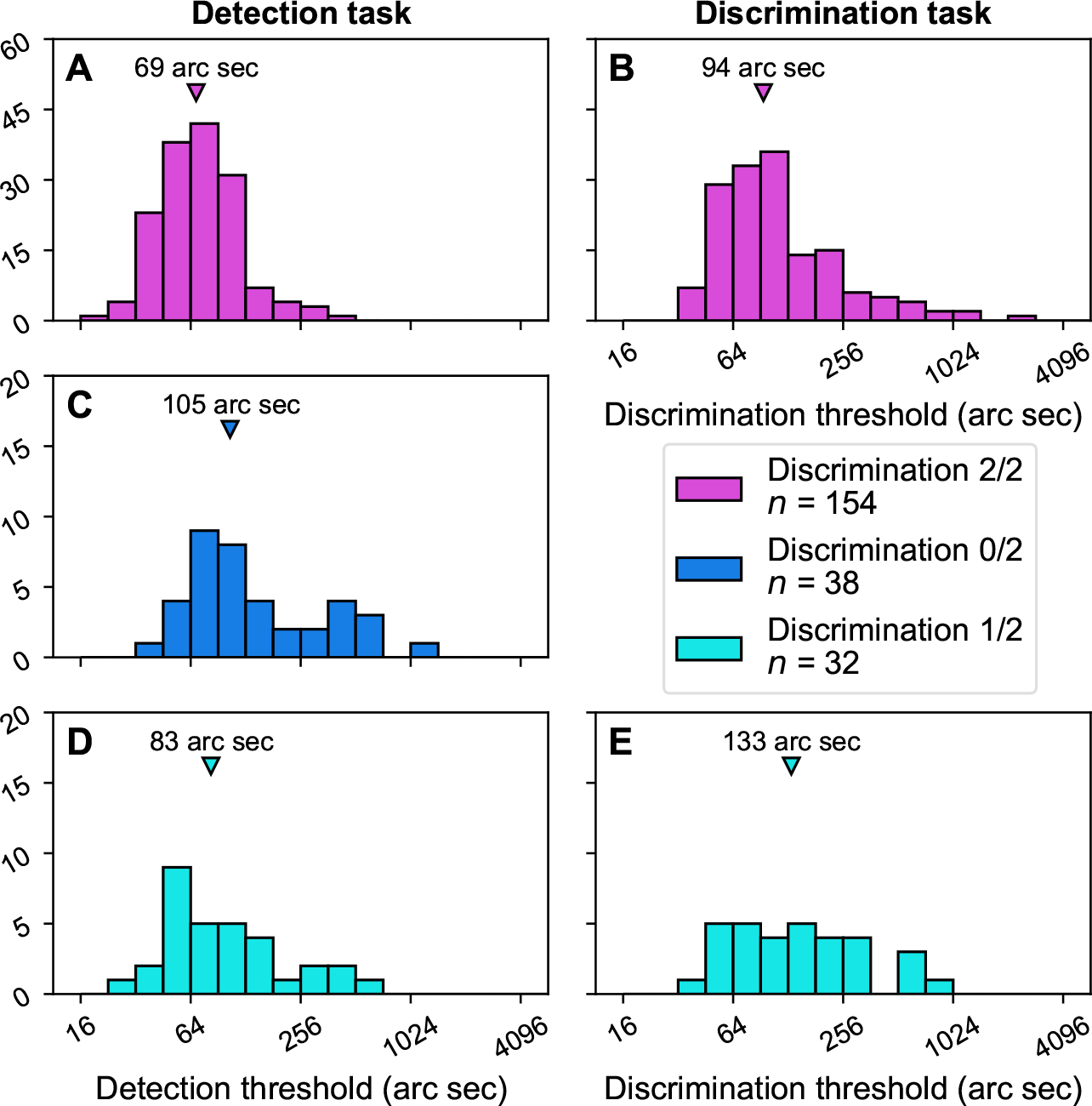
Histograms with distributions of detection (first column) and discrimination (second column) thresholds obtained from the three groups which are divided by the number of valid results obtained on the discrimination task (as in Figure 3). The triangle marker indicates the median of each distribution.

A Kruskal-Wallis H-test was performed on the detection threshold distributions, finding significant variation between them (*H*(2) = 26.0, *P* < 0.001). Pairwise comparisons made using Mann-Whitney U tests found a highly significant difference between detection thresholds from the groups who had either two or zero valid discrimination threshold measurements (*U* = 1404, *n*_1_ = 154, *n*_2_ = 38, *P* < 0.001). Analyses made with the group who had one valid discrimination threshold showed a significant difference compared to those with either two (*U* = 1897, *n*_1_ = 154, *n*_2_ = 32, *P* = 0.041) or zero (*U* = 790, *n*_1_ = 38, *n*_2_ = 32, *P* = 0.032) valid thresholds.

For the discrimination task (right column of Figure 4), the median threshold was lower in the group who obtained valid results on both repetitions (94 arc sec) compared to the group who only obtained a valid result on one repetition (133 arc sec). Comparing discrimination threshold distributions between these groups, we find again that the difference is significant (*U* = 1916, *n*_1_ = 154, *n*_2_ = 32, *P* = 0.048).

Finally, we compared detection thresholds against discrimination thresholds from the two groups (top and bottom rows of Figure 4). In both groups, the median discrimination threshold was higher than the detection threshold. For participants who obtained two valid measurements on both tasks, the difference in performance between the two was highly significant (Wilcoxon signed-rank test *Z* = 1413, *n* = 154, *P* < 0.001). For participants with only one valid discrimination measurement, the difference was significant (*Z* = 145, *n* = 32, *P* = 0.025).

## 4 Discussion

Previous investigations of the prevalance of stereo-anomaly have found inconsistent results, ranging from 1-2% (Newhouse and Uttal, 1982; Birch et al., 2008; Bohr and Read, 2013) to 30% (Richards, 1970; Hess et al., 2015, 2016). These previous studies have differed in the methods used to measure stereoacuity. In this study we compared the incidence of stereoacuity measured using two tests with similar stimuli but different task design in the same population.

We found that 98% of our participants gave valid results in both repetitions of a disparity detection task. The remaining 1-2% proportion of stereo-anomalous participants agrees with the previous studies that found lower prevalence of the condition (Newhouse and Uttal, 1982; Birch et al., 2008; Bohr and Read, 2013). Within the group that could perform the detection task however, only 69% were able to perform both repetitions of an otherwise equivalent discrimination task. Of the other 31%, there were 14% who were able to perform one of the two repetitions. The remaining 17% were unable to perform either repetition of the discrimination task. We therefore propose that the more common stereo-anomaly concerns the identification of the sign of disparity. The 30% value from the previous literature may reflect a combination of individuals who have difficulty with discrimination tasks (our 14%) and those who are simply unable (our 17%). Our results indicate that individuals who have difficulty with the discrimination task perform worse at the detection task, compared to those who are able to perform disparity discrimination.

We performed additional analyses to investigate possible explanations for this result. Our analysis of the overall probability of responding correctly (regardless of disparity) found that the individuals we class as having difficulty with disparity discrimination tend to show above-chance performance (even when a valid result is not obtained). On the other hand, those we define as “unable” typically perform at chance when faced with a discrimination task. Although the majority of those with only one valid measurement performed better in the retest measurement compared to the test measurement, our Bland-Altman analysis did not reveal a general test-retest bias for either the detection or discrimination task.

Results from other tasks can further narrow-down our interpretation. In our other studies using similar methods we did not find the large proportion of stereo-anomalous individuals identified in this study. In Alarcon Carrillo et al. (2020) we looked for, but did not find, stereo-blindness specific to detecting stimuli in one direction of disparity (Richards, 1971). We speculate that a deficit of this kind would also give a different result in the current study, as the ability to see one direction of disparity should still allow participants to perform the discrimination task. Similarly, Alarcon Carrillo et al. (2023) introduced an “odd-one-out” version of the test where there were four wedges. Three wedges had the same disparity (crossed or uncrossed), the target was the fourth wedge, which had the opposite sign of disparity. Among the 17 control participants in that study, we also did not find any stereo-anomalous participants. This suggests that identifying the direction of disparity may introduce an additional difficulty for some participants which is not present in tasks which only require them to discriminate an odd-one-out.

Specific difficulties related to the discrimination of disparity information have been noted previously (Harris et al., 2012). It has been suggested that sensitivity in depth discrimination tasks relates to the past stereoscopic viewing experience of the participant (Stransky et al., 2014). Perhaps conversely, a negative association with age has also been found. In discrimination experiments using stimuli with both crossed and uncrossed targets (similar in that way to our design), better performance was found in children aged 4-6 years old compared to adults (who were at chance; Wilcox et al., 2017). We expect the portable tablet-based testing approach used in the current study will provide an efficient means of identifying the possible stimulus and task factors that show this stereo-anomaly in future work.

## 5 Additional information

## 5.1 Acknowledgements

The authors would like to thank Ran Zhang for assisting in data collection. This work was supported by the Natural Sciences and Engineering Research Council of Canada (NSERC grant to RFH #2016-03740), and the Natural Science Foundation for Distinguished Young Scholars of Zhejiang Province, China (Grant No. LR22H120001 to JZ).This pre-print manuscript was written in L^A^TEX using TeXShop, and is formatted with a custom style available at: github.com/alexsbaldwin/biorxiv-inspired-latex-style.

## 5.2 Author contributions in CREDIT format

**Alex S Baldwin**: Conceptualization, Methodology, Software, Validation, Formal Analysis, Investigation, Data Curation, Writing - Original Draft, Writing - Review & Editing, Visualization, Supervision, Funding Acquisition; **Seung Hyun Min**: Investigation, Writing - Review & Editing; **Sara Alarcon Carrillo**: Formal Analysis, Writing - Review & Editing; **Zili Wang**: Investigation, Writing - Review & Editing; **Ziyun Chen**: Investigation, Writing - Review & Editing; **Jiawei Zhou**: Resources, Writing - Review & Editing, Supervision, Project Administration, Funding Acquisition; and **Robert F Hess**: Conceptualization, Methodology, Resources, Writing - Review & Editing, Supervision, Project Administration, Funding Acquisition.

## 5.3 Potential Conflicts of Interest

Alex S. Baldwin (ASB) and Robert F. Hess (RFH) are both inventors on patent(s) and other intellectual property (including Hess and Baldwin, 2020) concerning stereovision and the measurement and treatment of disorders of binocular vision such as amblyopia. Some of these technologies have been commercially licensed by McGill University to Novartis International AG. The inventors have (separate from the work described here) worked to develop these technologies for clinical use through a research agreement involving Novartis and the Research Institute of the McGill University Health Centre.

## 6 Appendix

The distributions of disparity thresholds found with the two tasks is shown in Figure A1. The two tasks are represented in the two rows (top row detection, bottom row discrimination). The two columns show data from the test (first column) and retest (second column) measurements.

**Figure A1:**
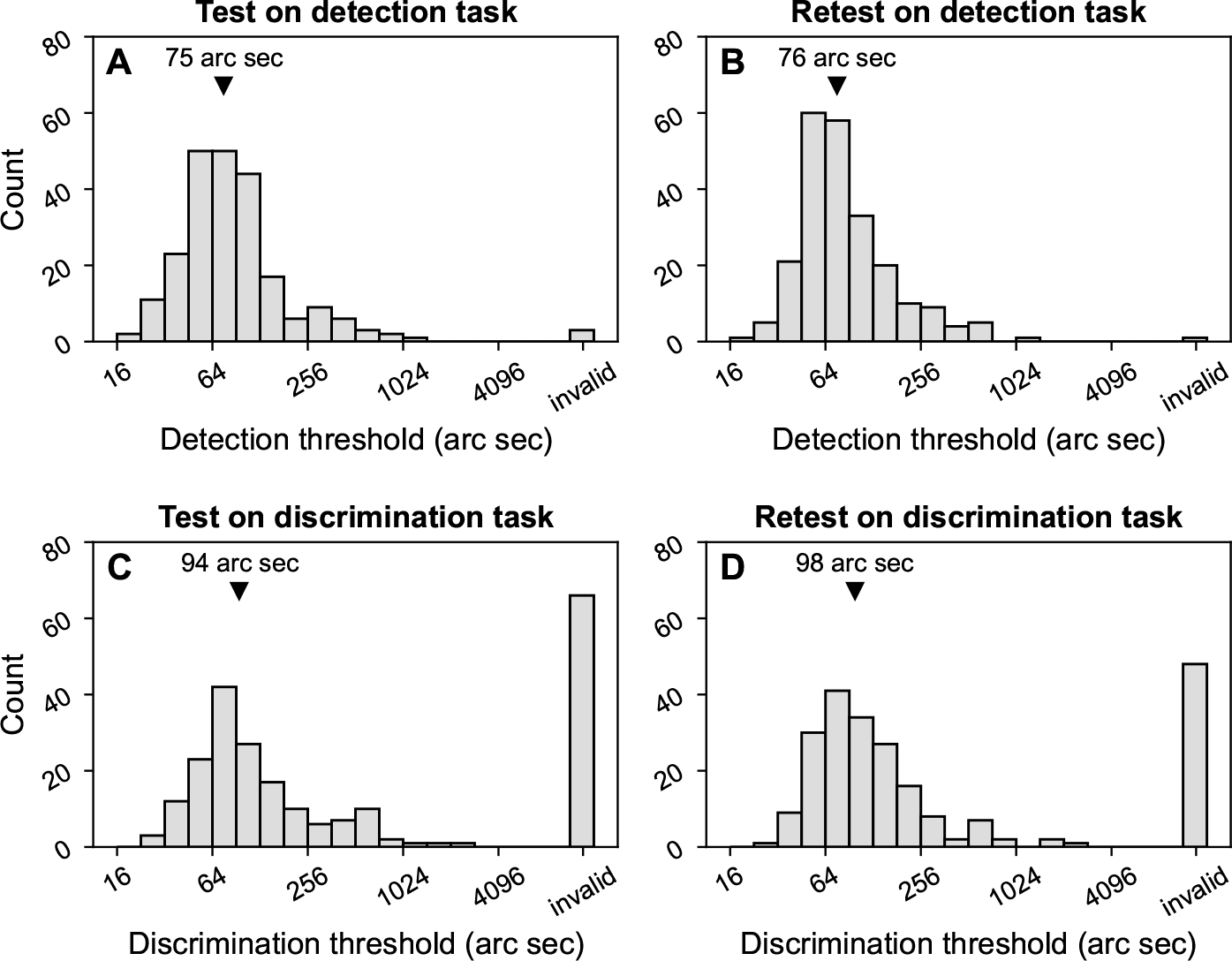
Histograms of disparity thresholds found with the detection (**A**-**B**) and discrimination (**C**-**D**) tasks. The subplots show data divided into test (first column) and retest (second column) measurements. Invalid measurements (see Methods) are represented at an arbitrary high value (14,000 arc sec). The median of each distribution (excluding invalid measurements) is indicated with a triangle symbol. Thresholds are presented from 228 participants per histogram.

Considering only the valid results, the median stereoacuity is higher in the discrimination task than the detection task for both the test (94 vs. 75 arc sec) and retest (98 vs. 76 arc sec) measurement distributions. All four distributions show some skewness. The Fisher-Pearson coefficient of skewness is 1.0 for both detection and discrimination in the test histograms. The retest histograms are more skewed. For the detection task the skewness coefficient is 1.1, for the discrimination plots the skewness coefficient is 1.2. Due to this skewness, the distributions were compared with Mann–Whitney U tests. There was a significant difference between the two distributions obtained in both the test (*U* = 13903, *n*_1_ = 225, *n*_2_ = 162, *P* < 0.001) and retest (*U* = 15835, *n*_1_ = 227, *n*_2_ = 180, *P* < 0.001) measurements. An analysis of the difference between test and retest measurements obtained with the two tasks was performed to look for any consistent bias. Bland-Altman plots are shown in Figure A2. The bias was not significant for any group in either task (see Table A1).

**Figure A2:**
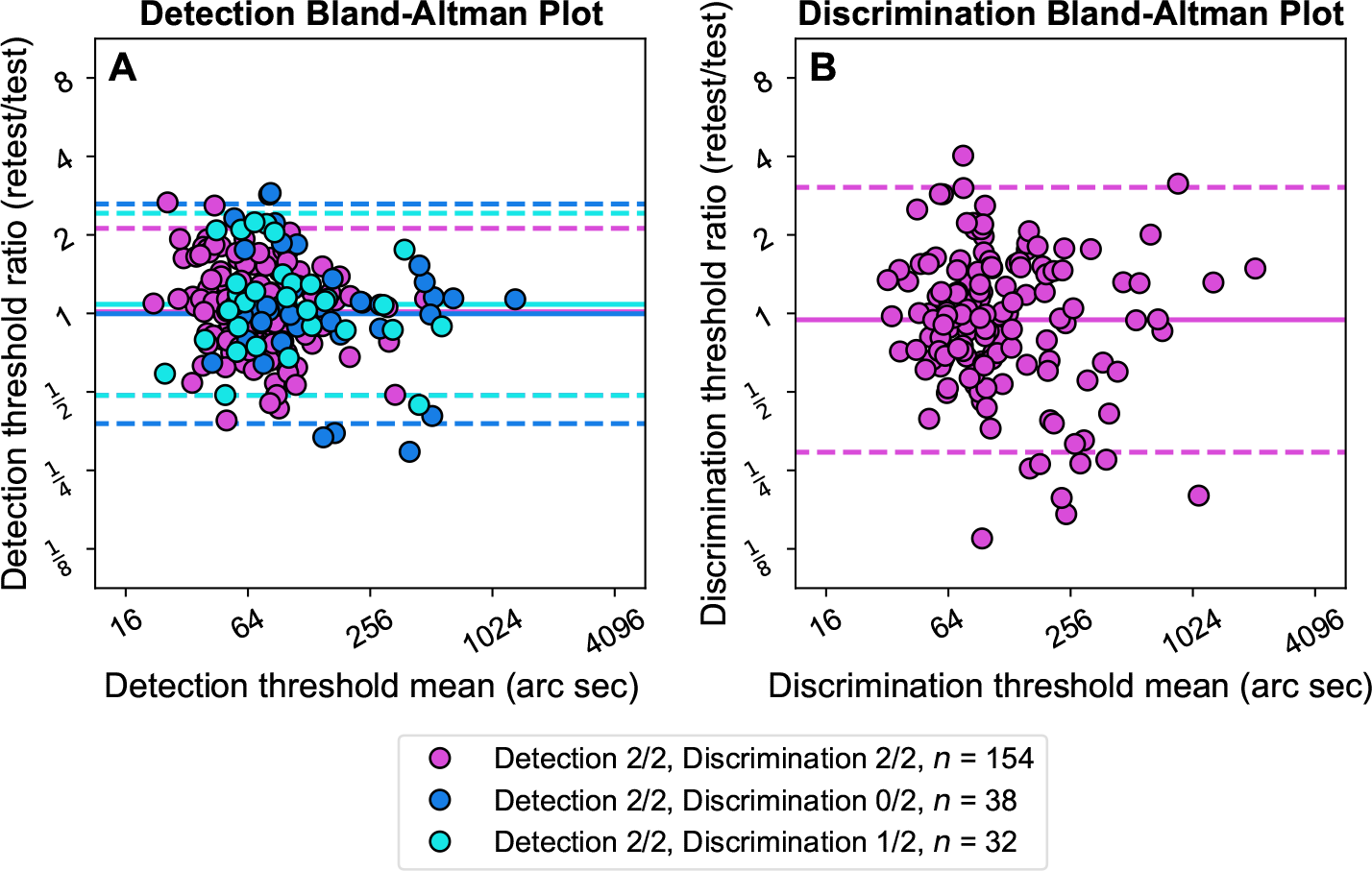
Bland-Altman plots showing relationships between the means of the test and retest thresholds for the detection (**A**) and discrimination (**B**) tasks, and the difference (of the log_2_ thresholds, so a ratio in linear units) between test and retest in each case. The solid horizontal line for each group shows the mean difference (the bias). The dashed lines show the *±*1.96 standard deviation range around that bias.

**Table A1:** Analysis of bias from Bland-Altman plots shown in Figure A2. Data are subdivided based on the number of valid results obtained with each task.

| Task | Group | Bias | One-sample t-test |
| --- | --- | --- | --- |
| Detection | Detection 2/2<br>Discrimination 2/2 | +0.02 | $t(153) = 0.45, P = 0.65$ |
| | Detection 2/2<br>Discrimination 0/2 | -0.01 | $t(37) = -0.04, P = 0.97$ |
| | Detection 2/2<br>Discrimination 1/2 | +0.12 | $t(31) = 1.09, P = 0.29$ |
| Discrimination | Detection 2/2<br>Discrimination 2/2 | -0.08 | $t(153) = -1.17, P = 0.24$ |

Figure A3 shows the overall proportion-correct data from the participants who performed poorly on one or both repetitions of the discrimination task. The data are divided based on whether an invalid result was found on both measurements, or only on the test or the retest measurement. For the participants from whom invalid results were obtained on both repetitions, half had overall P(correct) performance that was not significantly different from chance on both test and retest. These 19 data points are not plotted on the graph. The remaining participants show above or below chance performance on either the test or retest. The participants who obtained a valid measurement on either the test or retest measurement tend to show above-chance performance overall.

The distribution of above- and below-chance performance obtained on the two repetitions is visualised in Figure A4. This is based on the same data as Figure A3, plotted in three bins per repetition. These indicate whether participants’ performance did not significantly differ from guessing, or if it was significantly above-or below-chance. Significance judgements were made on the basis of the 99% binomial confidence intervals.

**Figure A3:**
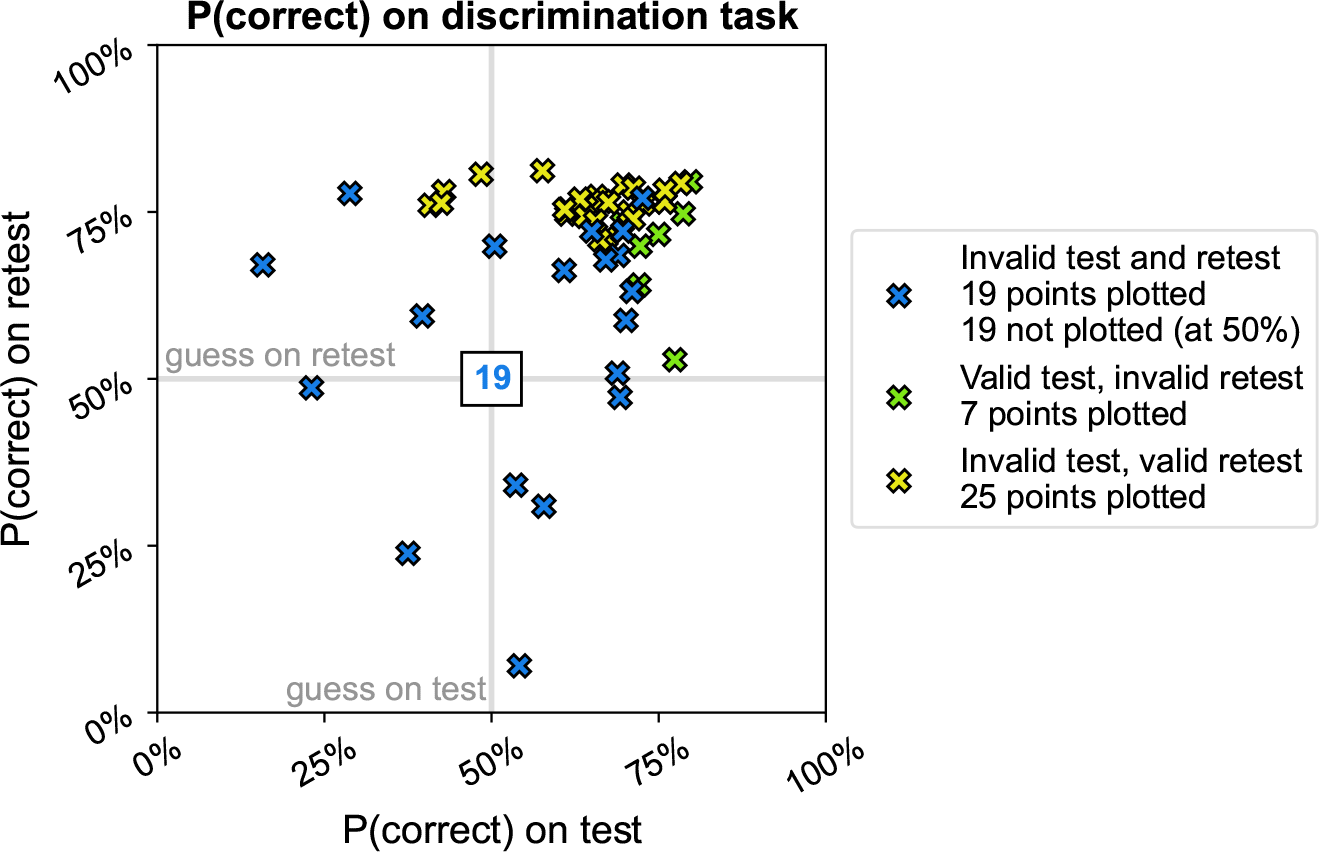
Overall proportion-correct data from the group of participants who performed poorly on the discrimination task. The overall P(correct) from the test and retest measurements are plotted on the x-axis and y-axis respectively. The guess-rate for the discrimination task is indicated by the grey lines at 50% on each axis. There were 19 data points omitted (indicated in the centre of the graph) based on the 99% binomial confidence intervals overlapping the guess rate for both the test and retest measurements.

**Figure A4:**
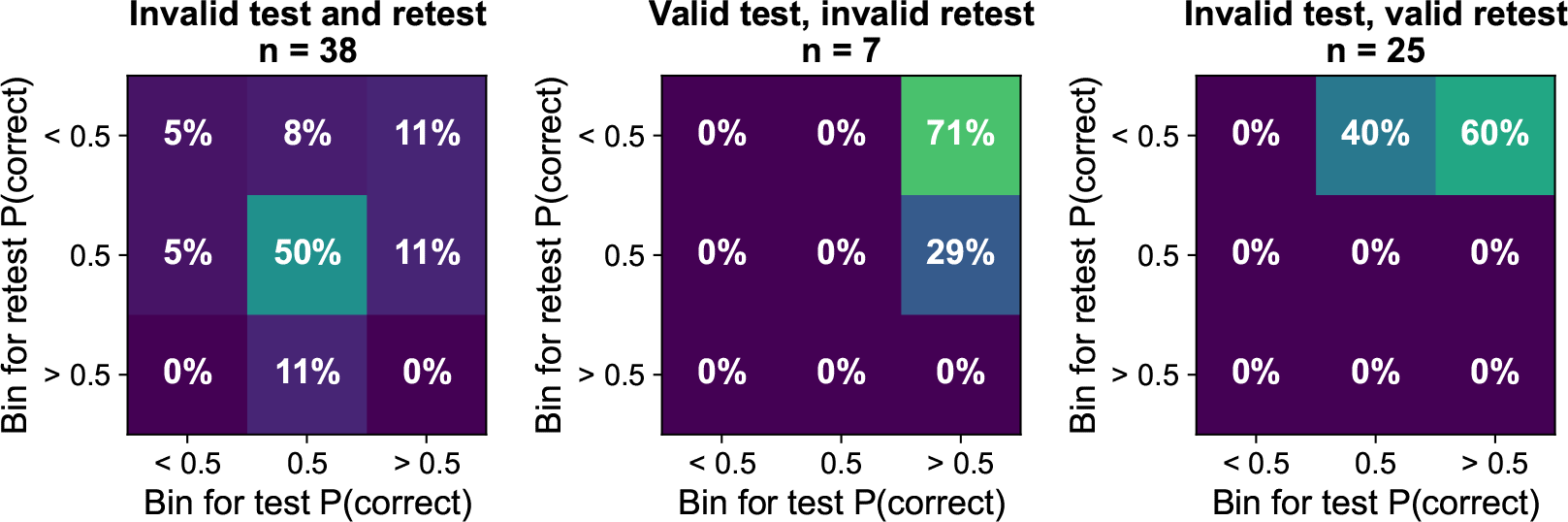
Data from Figure A3 binned according to whether participants performed significantly (by the 99% binomial confidence intervals) above or below chance (50%) on the test and retest measurements of the discrimination task.

No participant performed significantly below chance on both test and retest measurement. Had anyone done so, this could indicate a misunderstanding of the task instructions. As noted above, from the group who gave invalid results on both measurements (first column), the majority of individuals were at chance for both test and retest repetitions. The remaining 50% are distributed quite evenly around the other bins.

For the individuals who obtained one valid measurement (second and third column of Figure A4), the P(correct) performance on the other (invalid) repetition also tends to be above-chance. In those groups, it is also notable that we do not see any significantly below-chance behaviour. This indicates that the difference in performance on the valid and invalid repetitions is not due to them performing the task “backwards” (consistently selecting the wrong stimulus) in one repetition.

